# Heterogeneity of NOTCH3 activation mechanisms uncovers therapeutic potential for targeted therapies for CADASIL

**DOI:** 10.1101/2024.11.17.623993

**Authors:** Samira Hosseini-Alghaderi, Martin Baron

## Abstract

NOTCH3 is a developmental signalling receptor that regulates vascular smooth muscle cell (VSMC) proliferation, maintenance and phenotypic plasticity. Its misregulation is linked to both age-associated and inherited forms of small vessel disease and vascular dementia. CADASIL is a dominantly inherited condition, linked to recurrent stroke and vascular dementia, which is caused by mutations in NOTCH3. The mutations result in the accumulation of the extracellular domain (ECD) of NOTCH3 which has a toxic gain of function effect. Misregulated signalling may also play a role. ECD detachment is an obligatory step in NOTCH3 activation, but the mechanisms of CADASIL mutant NOTCH3 activation are not understood. Some CADASIL mutants affect ligand-induced activation and so ligand interactions are not a common underlying requirement for disease progression. Here we show that both wild type and CADASIL mutant NOTCH3 proteins are endocytosed as a full-length protein and then undergo dissociation and independent trafficking of the ECD and ICD in the endosome. We further show heterogeneity of activation mechanisms between different CADASIL mutants defined by dependency or not on metalloproteases. Understanding the variety of mechanisms by which NOTCH3 signalling and ECD shedding occur will inform new targeted approaches to treatments of small vessel disease. Tuning NOTCH3 activity through modulation of the endocytic pathway may offer better tolerated approaches than direct targeting of NOTCH3 signalling, important when considering the balance of risk and benefit for treating long term, chronic, progressive diseases, like vascular dementia.

## Introduction

Small vessel disease (SVD) is an age-associated, progressive and incurable condition, linked to impaired vascular smooth muscle cells and an important contributor to stroke and vascular dementia, the latter comprising around 20% of all dementia diagnoses^1,2^. Around 50% of people aged over 65 have signs of SVD on autopsy^3^. The signalling receptor NOTCH3 is a key player in both age-associated and genetically inherited forms of SVD because of its role in regulating phenotypic plasticity and maturation of vascular smooth muscle cells (VSMCs)^3–12^. NOTCH3 is therefore an attractive target for developing therapies linked to VSMC misregulation. Indeed, there is already significant interest in targeting Notch proteins for other therapeutic applications including cancer^13^. However, direct targeting of Notch to switch off the pathway can have severe side effects^14^. The level of risk that is associated with such treatments is not appropriate for chronic, progressive diseases that are not immediately life threatening, making finding effective treatments problematic. Understanding how to manipulate NOTCH3 signalling in more subtle ways can provide novel routes towards new therapies.

Work on *Drosophila* Notch has shown that the endocytic pathway has the capacity to tune the level of Notch activity, both up and down, through modulating surface exposure of Notch to ligands, and regulating the activity of ligand-independent activation of Notch localised on the endosomal surface, to provide robustness to environmental and genetic perturbation^15,16^. NOTCH3, one of four human Notch genes, has also been demonstrated to undergo ligand-independent activation although the mechanism has not been explored^17,18^. NOTCH3 is involved in the regulation of vascular smooth muscle cell (VSMC) maturation, and certain missense mutations are linked to a dominantly inherited vascular dementia disease CADASIL^19,20^. The latter is characterised by accumulation of the detached NOTCH3 extracellular domain (ECD), which comprises of 33 tandem Epidermal Growth Factor (EGF)-modules (Figure 1A), a structural motif that is defined by three conserved disulphide bonds^21^. Archetypal CADASIL mutations, located in the extracellular EGF repeats, either add to, or remove, a cysteine residue. There is heterogeneity of disease outcomes reported depending on the EGF-module affected. EGF 1-6 mutations normally result in more severe and earlier-onset disease than mutations in other regions of the ECD^22^. CADASIL is associated with degeneration of vascular smooth muscle cells, but the underlying link to NOTCH3 misregulation is unclear, and may be multifactorial. Both altered NOTCH3 signalling and toxicity resulting from ECD accumulation in Granular Osmiophilic Material (GOM) deposits contribute to disease progression^23–28^. As with other Notch family proteins, NOTCH3 signalling is initiated by ligand binding, which results in proteolytic removal of the ECD by the metalloprotease ADAM10, and subsequently the release of intracellular domain (ICD) from the membrane by gamma-secretase^10^. The released ICD relocates to the nucleus to bind to and activate the CSL (CBF, Su(H), Lag2) transcription factor. Mutations that do or do not disrupt ligand binding and signal induction have been shown to cause CADASIL, indicating that altered ligand-dependent interactions are not a common underlying factor^23,29,30^. In *Drosophila melanogaster,* ligand-independent Notch activation is associated with endocytosis of full-length Notch and ectodomain shedding of the ECD within the endocytic pathway, as a precursor to gamma-secretase release of the intracellular domain from the endosomal membrane^15^. We therefore investigated the endocytic trafficking and processing of NOTCH3 and the consequences on these processes of differently located CADASIL mutants.

**Figure 1.**
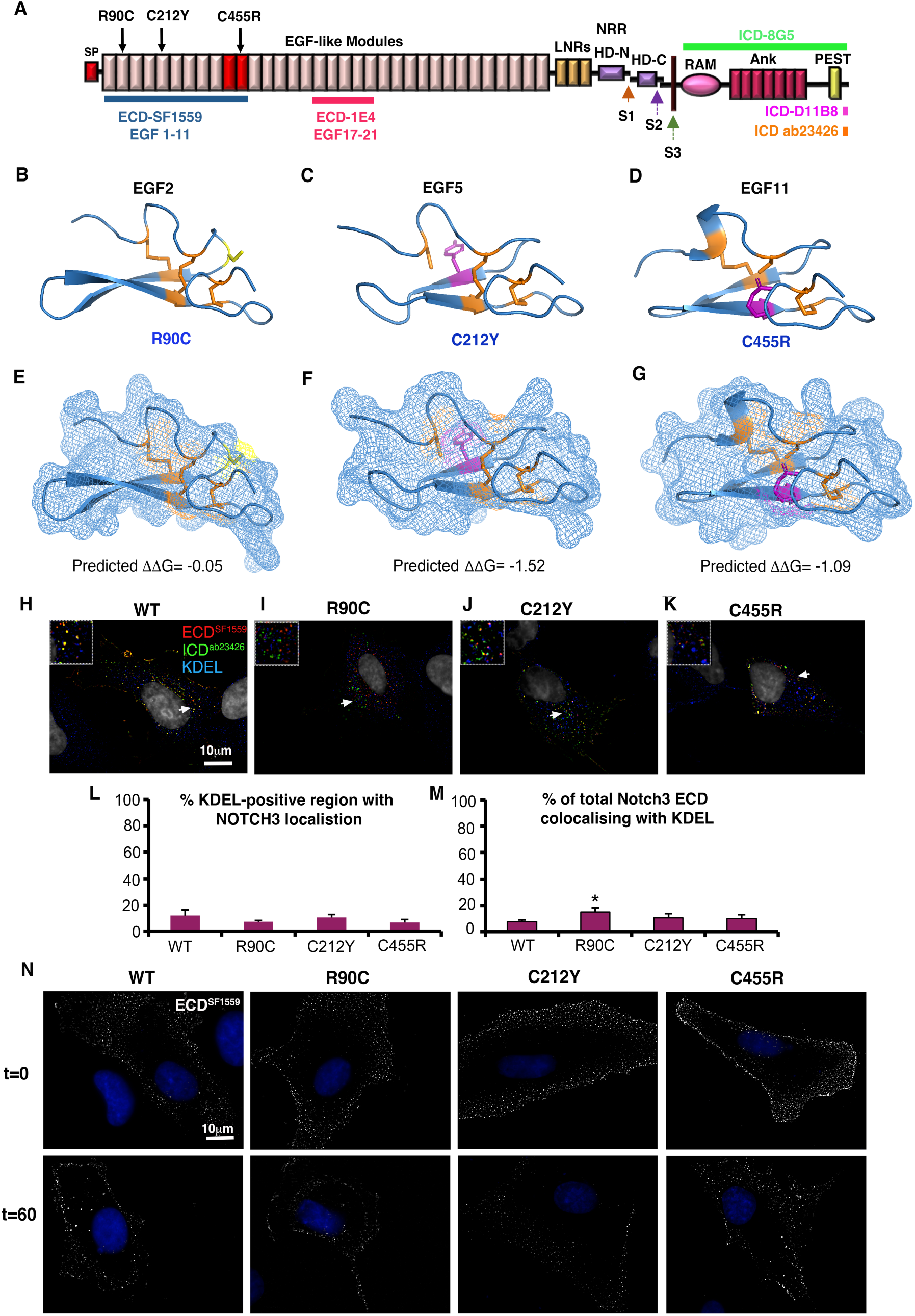
CADASIL NOTCH3 mutants are localised to cell surface like WT. **(A)** Modular structure of NOTCH3, SP indicates signal peptide, red shaded EGF-like modules indicate core ligand binding region, LNR indicates Lin12-Notch repeats, NRR is the negative regulatory region, S1, S2 and S3 are proteolytic cleavage sites, HD-N and HD-C are the heterodimer interface region. RAM and Ankyrin repeat region (Ank) are motifs involved in binding CSL transcription factors, PEST domain is involved in ubiquitin-dependent turnover of the ICD. Location of mutants used in this study are indicated by arrows. Coloured bars indicate locations of regions of NOTCH3 in which epitopes used in this study are located. **(B-D)** AlphaFold2 predictions of WT NOTCH3 EGF modules 2 **(B)**, 5 **(C)** and 11 **(D)**, and mutant substitutions introduced using the SDM programme. Location of the new Cys residue in R90C indicated in yellow, and other substitutions that replace as cysteine residues are shown in magenta. **(E-G)** Surface views of the structures from (B-D) depicted as a surface mesh. The introduced cysteine in R90C is surface exposed **(E)**, but the free cysteines in C212Y and C455R are buried within the module structure. ΔΔG values indicate predicted decreases in stability. **(H-K)** Expressed WT NOTCH3 **(H)** and CADASIL mutants **(I-K)** do not show strong accumulation in the ER when expressed in hTert-RPE1 cells. **(L, M)** scoring of NOTCH3 localisation in KDEL-positive ER, by % area of ER occupied **(L)** and by % of total NOTCH3 staining intensity **(K)** in z-sections through the cytoplasmic region. R90C shows a small increase in %NOTCH3 which is localised to the ER. * indicates P<0.05 by student t-test, error bars SEM, n=10. **(N)** Antibody surface labelling of NOTCH3 localisation on non-permeabilised cells at time 0 and 60 minutes during a live cell anti-ECD uptake assay. Similar time zero surface distributions, and subsequent surface depletion of WT and CADASIL mutants can be seen.

We found that both wildtype (WT) and CADASIL mutant NOTCH3 proteins traffic normally to the cell surface and display a punctate distribution on the cell membrane, from where NOTCH3 is endocytosed into the cell. In both WT and CADASIL mutants we found that the ICD becomes separated from the ECD during endocytosis and ICD subsequently localises independently of the ECD to the late endosome/lysosome. Expressed WT NOTCH3 undergoes a basal activation, independently of ligands but dependent on metalloprotease activity. Inhibition of the latter activity reduced ECD separation resulting in more full-length NOTCH3 in the late endosome. CADASIL mutants R90C, C212Y and C455R are also activated by this basal NOTCH3 activity. However, we found heterogeneity of activation mechanisms depending on the mutant tested. The R90C mutant switched its activation mechanism to a metalloprotease-independent and TRPML-dependent mechanism, while C212Y and C455R activated by a metalloprotease-dependent mechanism. This altered R90C mechanism was associated with differences in immunofluorescence localisation that suggest the ECD is detached before reaching the EEA1-positive endosome, unlike WT and the other CADASIL mutants tested. Different endocytic ectodomain shedding mechanisms and signal activation pathways may be relevant as a source of the long-term accumulation of the mutant NOTCH3 ECD in VSMCs and for understanding heterogenous pathological outcomes in CADASIL patients.

## Results

### Expressed CADASIL mutants are trafficked to the cell surface similar to WT

We studied the consequences on NOTCH3 of three CADASIL mutations, R90C, C212Y and C455R. R90C and C212Y mutations are located within the N-terminal EGF1-6 region where the majority of CADASIL mutations are located^4,31,32^. C455R was chosen as it is a mutation in the ligand-binding region of NOTCH3 which strongly inhibits ligand-dependent NOTCH3 activation^23,30^. The R90C mutation is in the top six of the most frequently occurring CADASIL mutants^33^ and C212Y is notable for spinal cord lesions in addition to typical CADASIL features^31^. The structural consequence of CADASIL mutations on the affected EGF domains of NOTCH3 have not been characterised in detail. It is largely assumed, based on HEK-293 cell overexpression studies, that such mutations disrupt the EGF-fold and this leads to accumulation of misfolded protein in the secretory pathway^34^. We used the structure prediction programme AlphaFold2^35,36^ to predict the structures of three wild type EGF modules from NOTCH3 in which CADASIL mutants have been previously described, EGF2, EGF6 and EGF12. We then used the Site Directed Mutator (SDM) programme^37^ to predict the consequences of three CADASIL mutations R90C, C212Y, and C455R) (Figure 1B-D). While a small decrease in stability was predicted for the three mutations (Figure 1E-G), the results showed that the missense mutations were accommodated with relatively little perturbation to the structure. Exposure of an unpaired cysteine residue on the surface of the EGF-module structure was only evident for the new cysteine introduced in the R90C substitution (Figure 1E-G).

When NOTCH3 was expressed in hTERT-RPE1 cells, previously used for similar studies of protein trafficking^38^, we did not observe strong accumulation of CADASIL mutant NOTCH3 in the endoplasmic reticulum (ER), compared to WT, that would indicate significant protein misfolding in the secretory pathway (Figure 1H-K). When we quantified ECD localisation we found no significant difference in the proportion of KDEL (K-Lysine, D-Aspartic acid, E-Glutamic acid, L-Leucine)-positive ER that was occupied by NOTCH3 immunolocalisation, although there was a small increase in the amount of R90C localised to ER measured by fluorescence intensity (Figure 1L, M). Surface staining of NOTCH3 localisation on live, non-permeabilised cells with an anti-ECD antibody showed a similar punctate surface distribution of WT and CADASIL mutant NOTCH3, which was depleted from the cell-surface over a one-hour chase period (Figure 1N).

### ECD separation after endocytosis of full-length NOTCH3

To investigate the fate of internalised NOTCH3, the uptake experiment was repeated and cells were further permeabilised and immunostained with anti-ICD and appropriate secondary antibodies after different chase intervals (Figure 2A-C). Up to the 15-minute chase time point we only observed ICD localised in EEA1 (Early Endosome Antigen 1)-positive early endosomes, consistent with lack of access of the anti-ECD to intracellular compartments during the pulse-labelling period (Figure 2D, E). Notch ICD was found to be localised in puncta on the EEA1-positive organelle. Notch ECD staining appeared in the organelle at 30 and 60-minutes time points, either colocalised with ICD as full-length Notch, or was present without ICD staining (Figure 2F, G), the latter suggesting separation of ECD from ICD had occurred. ECD of the three CADASIL mutants was also found localised within the early endosome after endocytic uptake and 60 mins chase (Figure 2 H-J). Therefore, these mutants follow a similar endocytic uptake to WT Notch3, even when the ligand-binding site is perturbed by the C455R mutant. The pulse chase experiment was repeated with a one-hour chase to investigate the localisation of WT NOTCH3 within CD63-positive late endosomes (Figure 2K, L). After a 1hour chase period, late endosomes were predominantly stained with anti-ICD, again in punctate subdomains within the CD63-positive structures (Figure 2L). There was little ECD colocalised with CD63, but outside of these late endosome structures both full-length NOTCH3 and separate ECD and ICD localisations were observed (Figure 2L).

**Figure 2.**
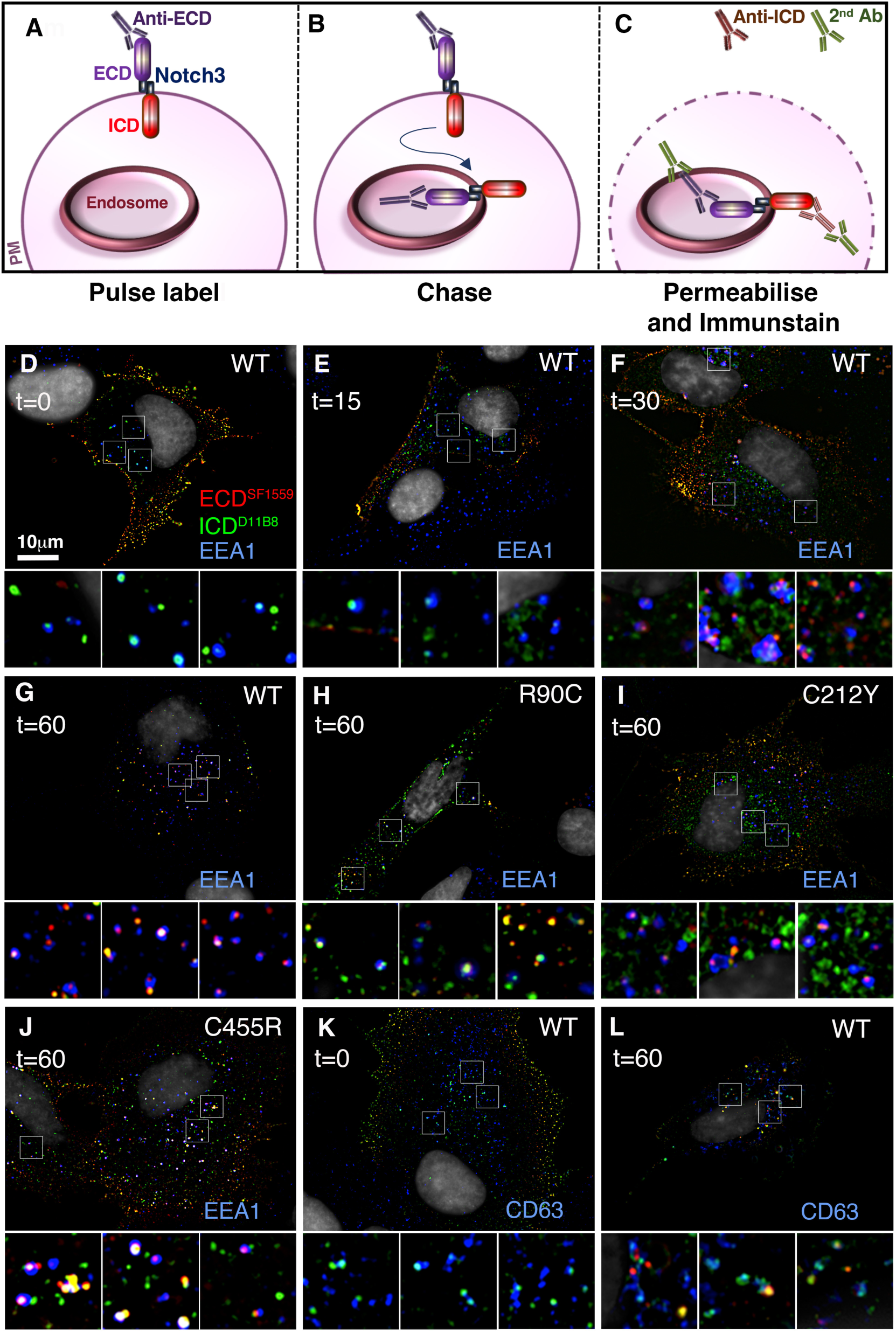
Full-length WT and CADASIL mutant NOTCH3 are endocytosed similarly. **(A-C)** Schematic view of pulse chase labelling protocol. **(D-G)** WT NOTCH3 localisation in permeabilised cells after antibody uptake at 0 **(D)**, 15 **(E)**, 30 **(F)**, and 60 **(G)** minutes. Little surface-labelled NOTCH3 is transported to the early endosome by 15 minutes chase but is evident in EEA1 positive endosomes by 30 and 60 minutes. Presence of both full-length and ECD-only staining suggests ECD-shedding can occur by the time NOTCH3 is localised to the EEA1-marked endosome. **(H-J)** CADASIL mutant NOTCH3 localisation after 60-minute chase shows the mutant proteins labelled at the cell surface also become localised to the EEA1-positive endosome. **(K, L)** After 60-minute chase surface, little labelled NOTCH3 ECD has reached the CD63-positive late endosome while ICD is present. However full-length NOTCH3 and ECD-only localisations are found adjacent to CD63-marked organelles.

These results indicate that surface-localised WT and CADASIL NOTCH3 is endocytosed as a full-length protein. The presence of ECD-only stained puncta within the early endosome suggests that this may be a location where ECD shedding occurs. We could not, however, distinguish whether ICD-only puncta in the early and late endosomes represented full-length NOTCH3 which was inaccessible to anti-ECD during the antibody-labelling period, or were products of separation of ECD from ICD.

To investigate further the endosomal localisation of NOTCH3, we immunostained permeabilised and fixed cells with anti-ECD and ICD, along with compartment markers for early endosome (EEA1), late endosome (CD63), recycling endosome (Rab11) and lysosome (LAMP1) (Figure 3A-H). In the EEA1-positive early endosome, WT NOTCH3 was predominantly either full-length or ECD-only localisation (Figure 3A). Full-length or ECD-only staining was also located within or adjacent to Rab11 marked recycling endosomes (Figure 3F). In contrast, in the CD63-labelled compartments we found both full-length and ICD-only spots (Figure 3G,I). Similarly, in LAMP1 marked lysosomes we predominantly found anti-ICD staining (Figure 3H, J).

**Figure 3:**
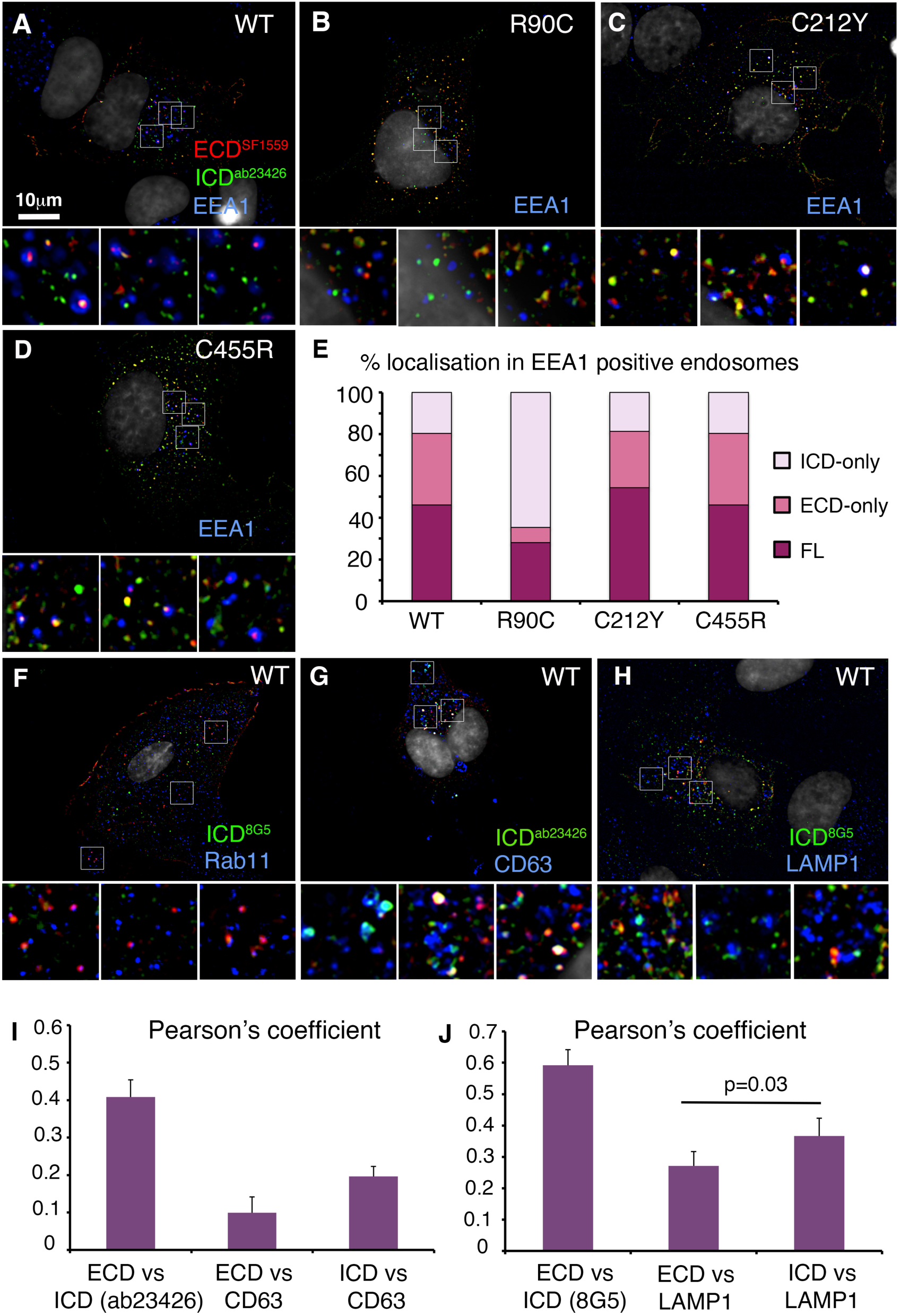
ECD-shedding in the early endocytic pathway. **(A-D)** Immunolocalisation of NOTCH3 with anti-ECD and anti-ICD compared with EEA1-marked endosomes in fixed and permeabilised cells of WT **(A)** and CADASIL mutants (**B-D)**. **(E)** Quantitation of NOTCH3 localisation in the early endosome, showing % endosomal-located NOTCH3 puncta which either full-length NOTCH3, ECD-only, or ICD-only staining. R90C is distinguished from WT and other mutants by a greater proportion of ICD-only puncta compared to ECD or full-length. For WT, n=91 NOTCH3 puncta from198 endosomes scored. For R90C, n=139 NOTCH3 puncta from 180 endosomes score. For C212Y, n=140 NOTCH3 puncta from 240 endosomes scored. For C455R, n=164 NOTCH3 puncta from 302 endosomes scored. Minimum of 7 cells scored for each construct. Data obtained from z sections from at least 7 cells. **(F)** Full-length NOTCH3, or ECD-only puncta located within or adjacent to Rab11-positive endosomes. **(G, H)** In CD63 positive endosomes **(G)** and LAMP1-positive lysosomes **(H)**, NOTCH3 localisation is predominantly as ICD-only puncta, quantified in **(I, J)**. Error bars in I and J are SEM, n=11 and n=7 respectively. Statistical significance by student t-test.

Interestingly, we found differences in the processing and localisation of the different CADASIL mutants in the early endosome. While C455R and C212Y mutants showed similar localisation of full-length and ECD-only puncta of NOTCH3 in the early endosome (Figure 3C, D, E), R90C showed a significantly higher proportion of ICD-only puncta and a decreased proportion of ECD-only spots (Figure 3B, E). Localisation of the NOTCH3 CADASIL mutants in the late endosome and lysosome was predominantly ICD-only immunostaining like WT (Supplemental Figure S1).

These results indicate that full-length NOTCH3 undergoes endocytic uptake and subsequent ECD removal, and that these processes are not affected by perturbation of the ligand binding site by the C455R mutation. There is a heterogeneity of outcomes arising from different CADASIL mutants, with the R90C mutation altering the location in which ICD localises independently from ECD. A common feature between WT and all three CADASIL mutants is the localisation of ICD staining in a punctate distribution in the late endosome and lysosome.

We considered whether differential epitope accessibility to anti-ECD and ICD antibodies might be a reason for observing separated ECD and ICD localisations. This may arise by epitope masking, partial fragmentation to release an epitope, or non-specific staining of either or both antibodies (Figure 4A). To address this, we conducted immunostaining with two different ECD antibodies raised against different regions of the protein with a single ICD antibody and *vice versa*. We found that when two ECD antibodies were used (SF1559 and 1E4) and ICD-only (D11B8) spots were still detected, lacking staining by both ECD antibodies (Figure 4B, D) indicating that epitope masking of the ECD is unlikely to account for observation of separate ICD staining. Furthermore, the colocalisation between the two ECD epitopes was significantly greater than the colocalisation between either ECD epitope and ICD as would be expected if the ECD and ICD were separating. These data therefore rule out significant epitope masking or proteolytic removal of just the N-terminal epitope. When we stained NOTCH3 expressing cells with two different ICD antibodies (D11B8 and 8G5) and a single ECD antibody (1E4), then ECD-only stained spots were still present lacking both ICD epitopes (Figure 4C, E), indicating that ICD epitope masking was unlikely to account for observation of separated ECD. Both ICD epitopes, but not the ECD epitope, were also detectable in the nucleus of NOTCH3 transfected cells (Figure 4 F-H). However, the colocalisation of the two ICD antibodies was not greater than the colocalisation of either epitope with the ECD and we saw less colocalisation of 8G5 epitope with the ECD than D11B8 with the ECD (Figure 4E). This suggested epitope masking of 8G5 epitope was occurring. To investigate this further we examined 8G5 localisation in the early endosome. Interestingly we found that unlike D11B8, 8G5 was rarely detectable in the early endosome or at the cell membrane (Supplemental Figure S2), while it was clearly present in the lysosome (Figure 3H, Supplemental Figure S1D-F). Epitope masking therefore seems likely for 8G5 epitope in the early endosomal trafficking pathway. However, we were able to rule this explanation out for D11B8 and ab23426 epitopes. All the antibodies stained transfected cells at a level well above any background staining in non-transfected cells, ruling out off-target immunostainings as an explanation for apparent epitope separation of expressed NOTCH3.

**Figure 4.**
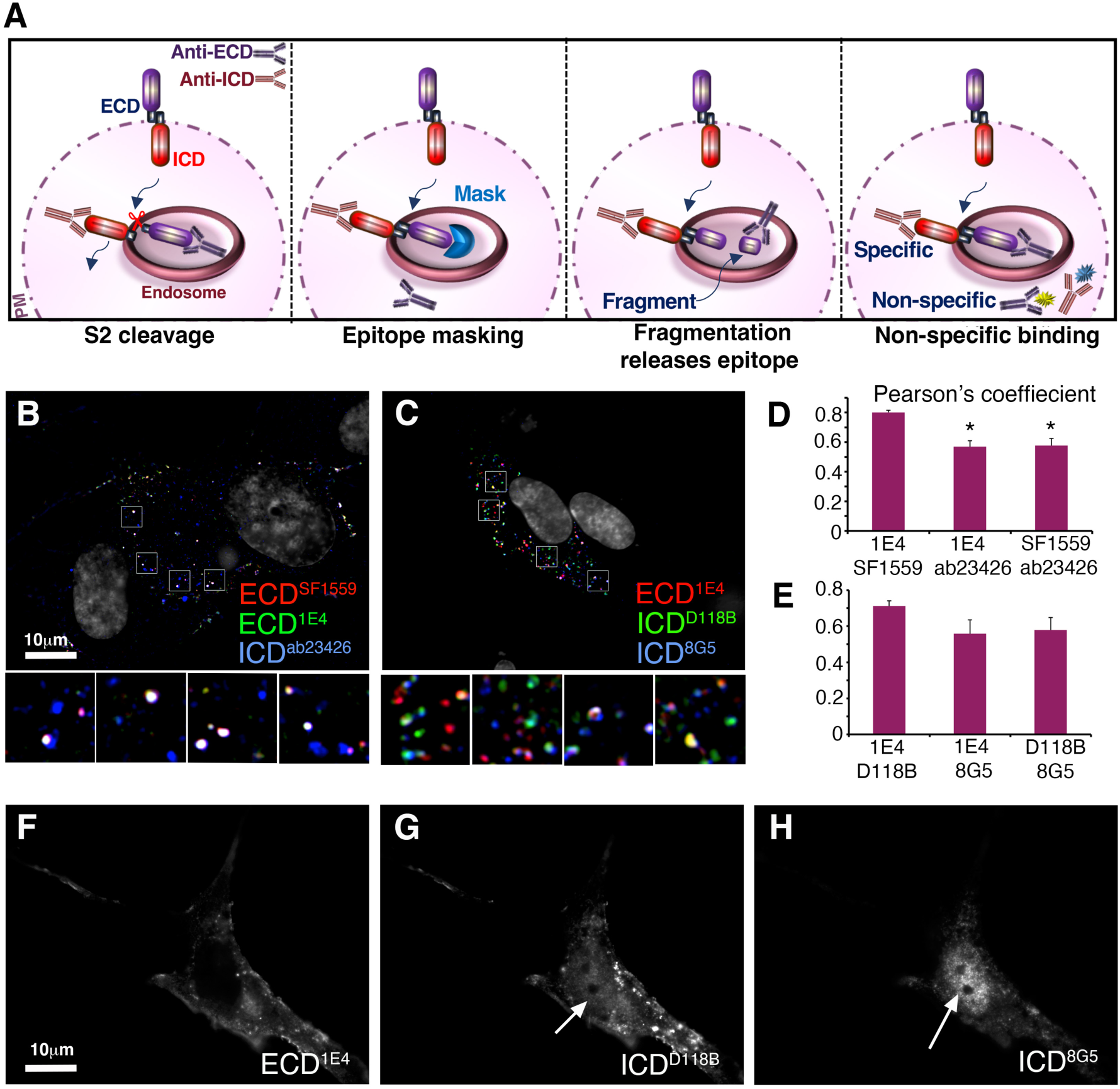
Visualisation of ECD/ICD independent localisation is not due to epitope masking. **(A)** Schematic diagram illustrating alternative explanations for visualisation of separated ECD and ICD-positive puncta, including: ECD shedding by S2 cleavage; epitope masking; fragmentation to remove terminal epitopes; non-specific staining. **(B)** Colocalisation of two different ECD epitopes compared to ICD^ab23426^. ICD-stained puncta can be observed which lack both ECD epitopes. **(C)** Colocalisation of two different ICD epitopes compared to anti-ECD^1E4^. ECD-only spots are present which lack both ICD epitopes. **(D, E)** Quantification of the colocalisation between the epitopes used in B, C. In D * indicates P<0.05 compared with double ECD epitope colocalisation by student t-test. Error bars represent SEM, n=5 **(D)**, n=14 **(E)**. **(F-H)** Both ICD epitopes are located to the nucleus in NOTCH3 transfected cells but not the ECD epitope.

We also utilised fluorescent protein tagged NOTCH3 constructs (Supplemental Figure S3A, B) to provide an alternative means to label the ECD and ICD, independent of antibody staining. FP1-NOTCH3 had mGFP-tagged ICD and mTagBFP2 located in the ECD (not visible in fixed samples). Non-permeabilised cells were stained with anti-NOTCH3 ECD. Colocalisation of anti-ECD immunostaining and GFP fluorescence confirmed that NOTCH3 located at the plasma membrane was full-length protein (Supplemental Figure S3C). In the FP2-NOTCH3 construct, the FP tags were mScarlet-I for the ECD and mNeon-Green for the ICD, which were chosen for their fast folding and pH resistance properties^39,40^. Consistent with the results using immunostaining, the FP2-tagged NOTCH3 showed separation of the ECD and ICD when transfected into hTERT-RPE1 cells (Supplemental Figure S3D). Furthermore, after costaining of permeabilised cells expressing FP2-NOTCH3 with anti-ECD, mNeon-Green labelled ICD puncta could be observed that lacked both anti-ECD staining and mScarlet-I fluorescence (Supplemental Figure S3D). The ECD antibody and mScarlet-I fluorescence showed a good colocalisation in cytoplasmic puncta, but at the outer cell membrane FP-tag fluorescence was weak compared to antibody staining. Nevertheless, regions of anti-ECD and mScarlet-I co-staining could also be observed at the cell membrane along with mNeonGreen fluorescence (Supplemental Figure 3D). The combined results using FP-NOTCH3 and anti-Notch antibody labelling are therefore consistent with ECD shedding within the endosomal trafficking pathway.

### Endogenous NOTCH3 undergoes ECD separation in the endosomal pathway

We next investigated whether the observed ECD/ICD separation in the endosomal pathway was a feature that resulted from elevated levels of NOTCH3 expression after transfection. For this we used two cell types with endogenous NOTCH3, the human breast tumour cell line MCF7 and human VSMCs derived by differentiation of hESCs.

To test the specificity of anti-NOTCH3 antibodies for immunostaining at endogenous expression levels we generated a knockout of NOTCH3 in MCF7 cells using CRISPR/Cas9 and compared the NOTCH3 immunostaining with the parental MCF7 line. The results (Supplemental Figure S4) showed virtually no background staining on knockout cells for anti-ECD (SF1559). The two anti-ICD antibodies D11B8 and ab23426 used in this study displayed some visible background staining in the nucleus, but not in the cytoplasm. Therefore, all three antibodies can be considered specific for endogenous levels of NOTCH3 staining in cytoplasmic puncta but unreliable for quantification of nuclear localisation.

Immunostaining of wild type MCF7 cells showed colocalisation of ECD and ICD around the perimeter of the cell, whereas NOTCH3 localisation within the cytoplasm could be as full-length, ECD-only or ICD-only labelled puncta. We detected all these forms in EEA1-labelled early endosomes (Figure 5A), but ICD-only was more prominent in CD63 labelled late endosomes (Figure 5B). When we treated the MCF7 cells with the metalloprotease inhibitor BB94 there was an increased presence of ECD colocalised with CD63 compared to control cells and full-length NOTCH3 could be visualised to be colocalised with CD63 (Figure 5C, E, F). This is consistent with a role for ADAM10 in the shedding of ECD by S2 cleavage. In contrast, treatment of cells with the gamma-secretase inhibitor (DAPT) did not cause an increase in full-length NOTCH3 in the late endosomes (Figure 5D) but there was a small increase in ICD localisation in the late endosome, (Figure 5G), consistent with the role of gamma-secretase in intramembrane S3 cleavage, which occurs after ECD shedding.

**Figure 5.**
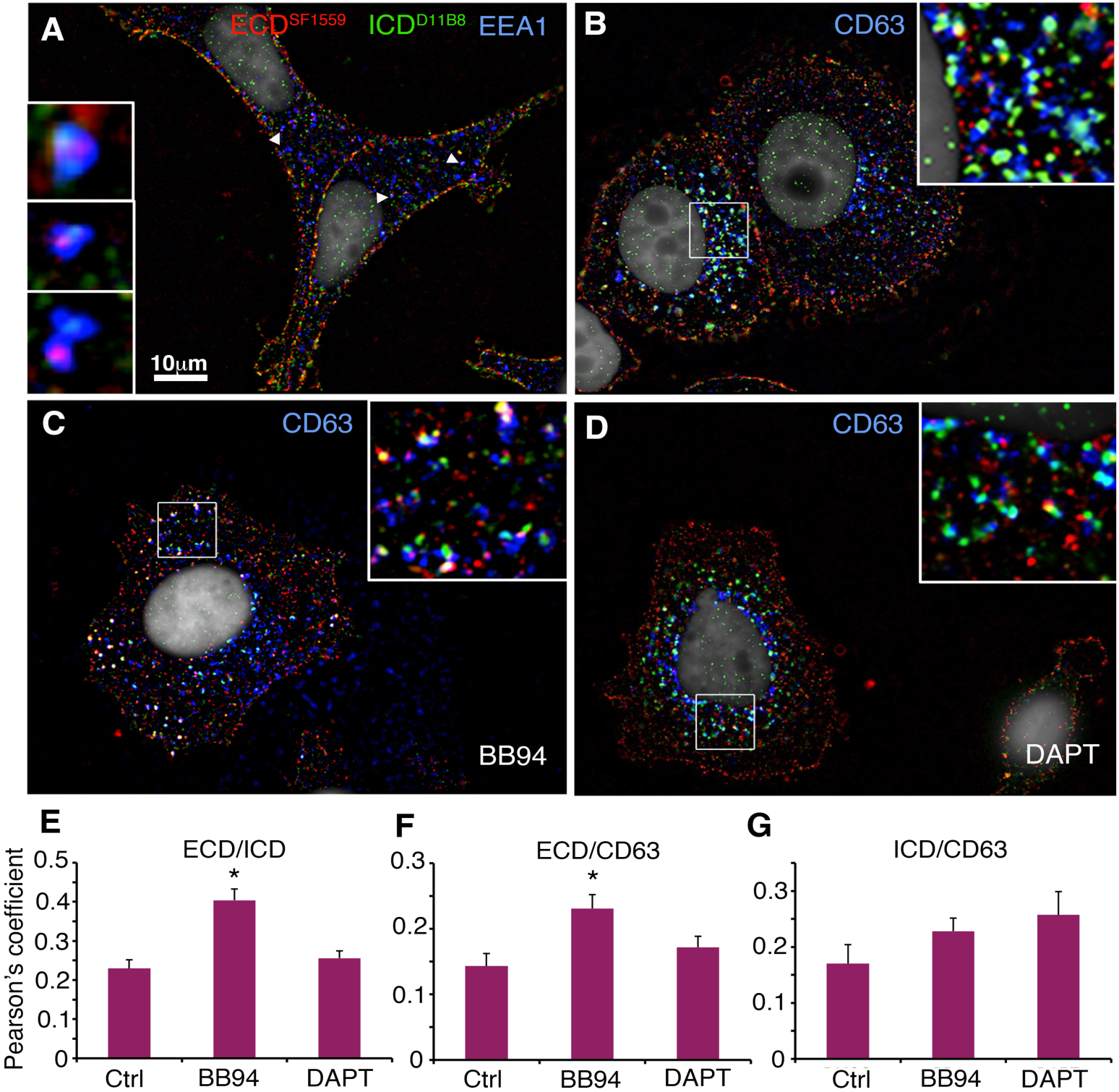
Effect of pathway inhibitors on endogenous NOTCH3 localisation in MCF7 cells. **(A)** EEA1-positive early endosomes of MCF7 cells costained with anti-ECD, and ICD. Arrowheads indicate endosomes shown enlarged in insets which have separate localisation of ECD and ICD puncta. **(B)** CD63-positive late endosomes costained with anti-ECD, and ICD. Boxed region is enlarged in inset showing late endosomes predominantly contain ICD-only puncta. **(C)** After treatment of MCF7 cells with metalloprotease inhibitor BB94, more full-length NOTCH3 staining is observed in the late endosomes, colocalised with CD63. **(D)** Treatment of cells with gamma-secretase inhibitor DAPT does not alter localisation of ECD in the late endosome. **(E-G)** Quantification of colocalization of ECD vs ICD **(E)**, ECD vs CD63 **(F)**, and ICD vs CD63 (**G)**. Error bars, SEM, control DMSO-only treated cells n=10, BB94 n=18, DAPT n=12. * indicates p<0.05 by student t-test compared to control.

Since the CADASIL mutant phenotype is manifest in VSMCs, we differentiated human embryonic stem cells into VSMCs using established protocols^41,42^ (Figure 6A-G), and confirmed differentiation by immunostaining for VSMC differentiation markers, Calponin and a-smooth muscle actin and change in morphology. We immunostained VSMCs for NOTCH3 ECD and ICD and costained for either EEA1 or CD63 (Figure 6H, I). We found that there was separate localisation of ECD and ICD immunofluorescence within the cells. EEA1 positive endosomes had mostly ECD-only localised in the central lumen of the endosome, but full-length or ICD-only spots were also present (Figure 6H). In CD63 positive late endosomes puncta of ICD-only spots were present but rarely any ECD-only puncta or full-length Notch3 (Figure 6I). These results are therefore consistent with our observations on over expressed NOTCH3.

**Figure 6.**
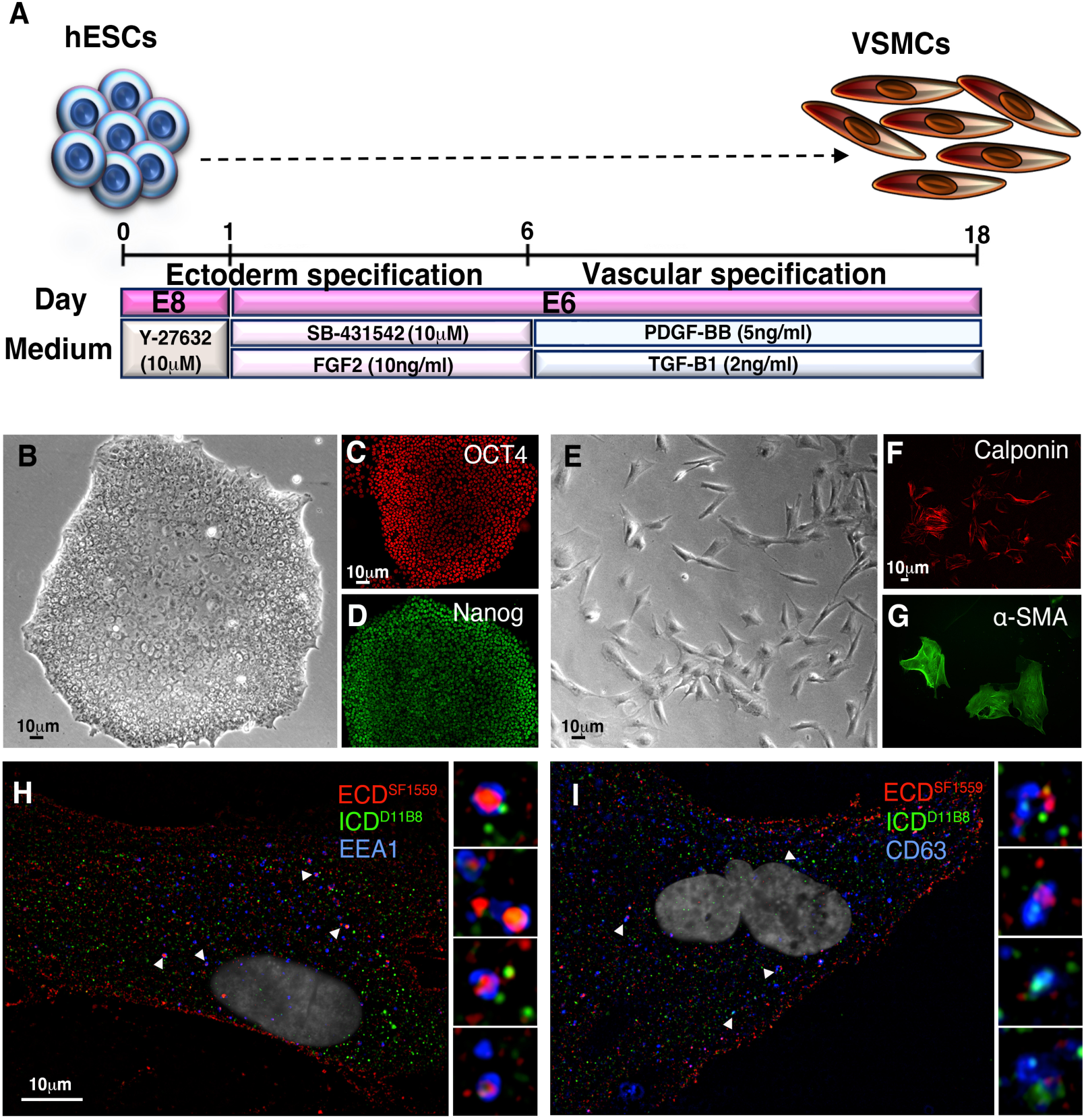
NOTCH3 ECD/ICD separation in endosomes of VSMCs derived from hESCs. **(A)** Summary of procedure followed to differentiate hESCs to VSMCs. **(B)** Brightfield image of undifferentiated hESC colony. **(C, D)** Undifferentiated hESC colony expressing stem cell markers OCT4 and Nanog. **(E)** Brightfield image of cells after differentiation protocol showing change of morphology. **(F, G)** After differentiation, cells express VSMC markers calponin and α-smooth muscle actin (SMA). **(H, I)** VSMCs immunostained to show separate localisations of ECD and ICD only puncta in early endosomes **(H)** and late endosomes **(I)**. Arrowheads point to endosomes shown enlarged in insets.

### Basal signal activation of WT and CADASIL mutant NOTCH3

We investigated whether NOTCH3 expressed in hTERT-RPE-1 cells was active in signalling. Cells were co-transfected with NOTCH3 expressing plasmid, a Notch response element (NRE) reporter driving luciferase expression, and a control plasmid that constitutively expresses Renilla for normalisation (Figure 7A). Relative signalling increased after transfection with WT NOTCH3 and this was reversed by treatment of cells with gamma-secretase and metalloprotease inhibitors (Figure 7B,C), but not by TRPML inhibitor. The latter affects the endolyosomal fusion/fission cycle^15,43^.

**Figure 7.**
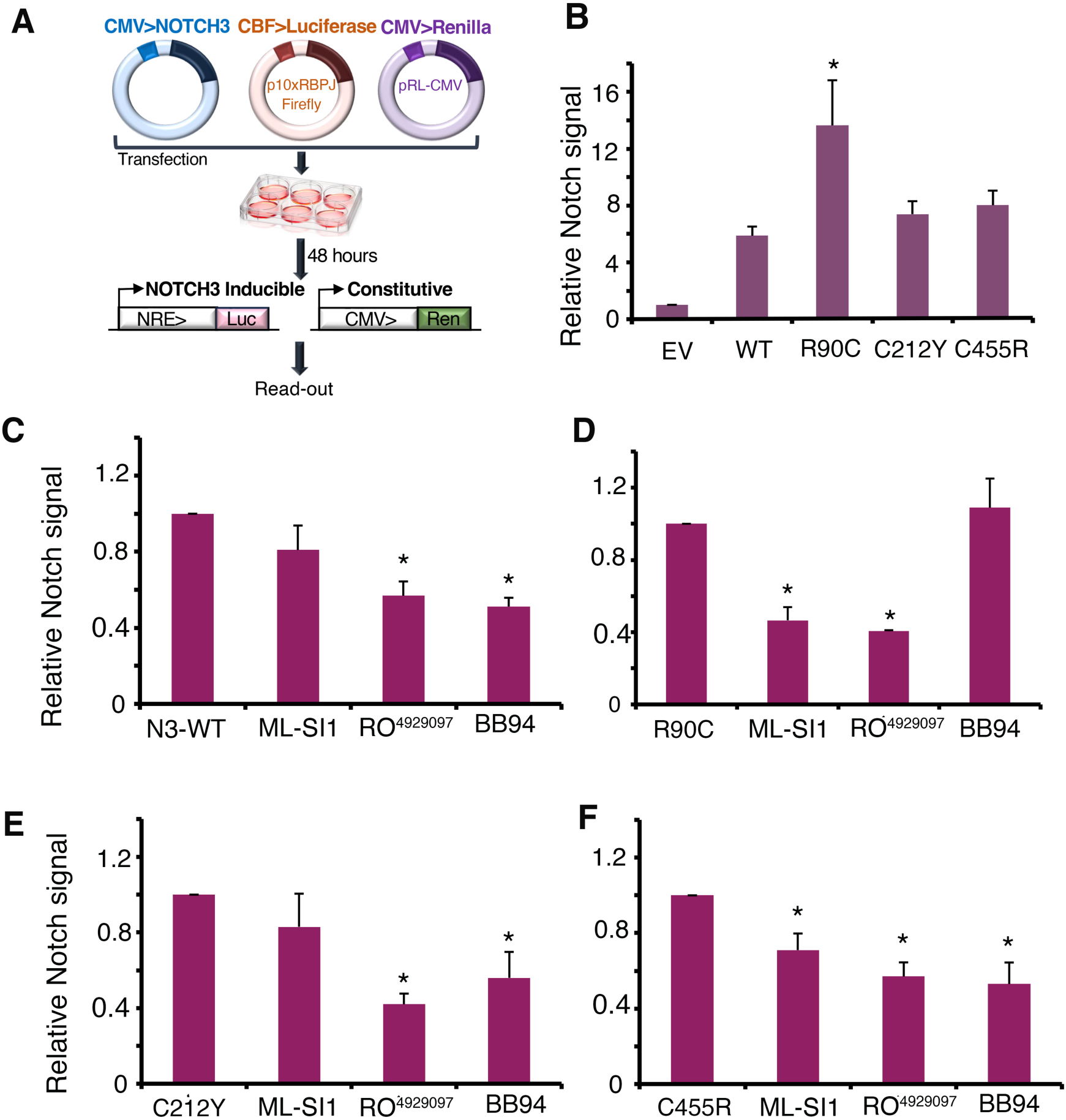
Basal NOTCH3 signalling requirements for WT and CADASIL mutants. **(A)** Schematic figure illustrating luciferase reporter assay methodology. **(B)** WT and CADASIL mutant NOTCH3 signalling after transfection into hTERT-RPE1 cells compared to empty vector (EV) control. * indicates p<0.05 compared to WT NOTCH3 by student t-test. Error bars SEM, minimum of n=4. **(C-F)** NOTCH3 signalling after treatment of transfected cells with TRPML inhibitor (ML-Sl1), gamma-secretase inhibitor (RO^4929097^), or metalloprotease inhibitor (BB94) for WT **(C)**, R90C **(D)**, C212Y **(E)**, and C455R **(F)** * indicates p<0.05 compared with control by student t-test. Error bars are SEM, minimum of n=3.

To investigate if CADASIL mutant NOTCH3 are also active for signalling, equivalent amounts of the CADASIL mutant constructs were transfected into hTERT-RPE1 cells. Basal level of signalling of C455R and C212Y was slightly elevated compared to WT but R90C showed significantly increased activity (Figure 7B). We next investigated whether signalling by CADASIL mutants involved the same or different mechanism compared to WT NOTCH3 using the panel of inhibitors (Figure 7C-F). Signalling by all constructs was reduced by gamma-secretase inhibitor. Unlike WT, R90C signalling was found to be metalloprotease-independent and but sensitive to TRPML inhibition. We found that C455R and C212Y were sensitive to metalloprotease inhibition and were less sensitive to TRMPL inhibition, hence these constructs signalled more like WT (Figure 7E,F). Thus, the increased signal activation of R90C mutation was associated with a switch in activation mechanism. A small but statistically significant decrease in signalling after TRPML that was observed for C212Y may suggests that the selection of activation mechanism is not all or nothing and both mechanisms may contribute to overall levels of activity for some mutants.

Our results therefore demonstrate heterogeneity of the cellular location where ECD shedding occurs and distinct basal activation mechanisms of different CADASIL mutants.

## Discussion

We investigated the endocytic trafficking and processing of human NOTCH3 and compared WT and three CADASIL mutant variants. We found that, contrary to expectations, the secretory pathway trafficking of mutant NOTCH3 was not blocked in the ER and both WT and mutant NOTCH3 were transported to the cell surface and displayed a similar punctate distribution. This indicates that protein misfolding is not a factor in CADASIL mutant outcomes. Modelling of mutant domains showed that amino acid changes could be incorporated without gross alterations to the EGF-fold.

Pulse chase endocytic uptake experiments showed that WT and mutant NOTCH3 were endocytosed similarly, becoming localised to the early endosome after 15-30 minutes of the chase period. Interestingly, we found both full-length (colocalised ECD and ICD staining) and ECD-only spots in the early endosomes, the latter normally located in the endosomal lumen. This indicates that, following uptake of full-length NOTCH3, ECD shedding occurs in the endocytic pathway. Even after a one hour chase we found little ECD in the late endosomes, although ICD was present. We could not distinguish whether this was due to ECD removal in the early endosome or because the endocytosed full-length NOTCH3 had not yet reached this location. However, when total NOTCH3 was immunostained in fixed/permeabilised cells, we also found a predominance of ICD-only staining in the late endosome and lysosome compared with ECD.

There are several reasons that immunofluorescence images could produce an apparent separate localisation of ECD and ICD epitopes even if NOTCH3 was present as full-length protein. We considered the possibilities of non-specific staining of either ECD or ICD antibodies, epitope masking, and a N or C-terminal truncation that results in near full-length NOTCH3 that lacks one of the terminal epitopes. All antibodies used in this study were specific to NOTCH3 expressing cells and not detectable in non-transfected cells at the exposures used, so all the staining observed could attributed to NOTCH3. To rule out epitope masking we used different antibody combinations to either double label the ECD or the ICD with differently located epitopes. We found ICD-only spots that lacked both ECD epitopes and we found ECD-only spots lacking both ICD epitopes. This supports the conclusion that true separation of ECD and ICD was the explanation for our imaging data since it is unlikely that both epitopes of the ECD or the ICD were masked at the same time. Only one epitope recognised by the rat monoclonal 8G5 antibody showed signs of epitope masking. While 8G5 stained the lysosome and nucleus of NOTCH3 expressing cells, it was barely detected in the early endosome or the cell membrane. This epitope exclusion was not observed for other anti-ICD antibodies used in this study. To further test our conclusions, we used double FP-tagged NOTCH3 as an alternative means to observe ECD/ICD separation that is not reliant on immunostaining and obtained results that support the finding that NOTCH3 is full-length at the cell membrane with ECD separation occurring in the endosome. The location of the ECD tag between the most C-terminal EGF modules and the NRR also implies that the majority of the ECD is removed together rather than N-terminal fragmentation being the cause of epitope separation.

We also investigated the possibility that the observed endocytic processing pathway was a result of over expressing NOTCH3. Two cell types were used, MCF7 which is a breast cancer cell line that expresses NOTCH3, and VSMCs differentiated from hESCs. The specificities of the antibodies used at these levels of endogenous expression were confirmed using a NOTCH3-knockout MCF7 cell line, generated by CRISPR/Cas9 induced non-homologous end joining. In both MCF7 cells and VSMCs we saw similar independent epitope localisation patterns as seen for over expressed cDNA in hTERT-RPE1 cells. Furthermore, when we treated MCF7 cells with the metalloprotease inhibitor BB94 we saw increased presence of ECD in the late endosome, colocalised with ICD and presumably representing increased trafficking of NOTCH3 full-length protein to the late endosome. This supports an intracellular location for ECD removal and demonstrates that the reduced ECD presence in the late endosome in untreated cells is not due to poor penetrance of the ECD antibodies into the late endosomal lumen.

To characterise the signalling activity for WT and CADASIL mutants we used a reporter assay that expresses luciferase in response to Notch pathway activation and treated cells with different pathway inhibitors. We found that expressed full-length NOTCH3 had basal activity that was dependent on metalloprotease and gamma-secretase activity. C455R and C212Y mutants showed a small increase in activity compared to WT, which was also reduced by the metalloprotease and gamma-secretase inhibitors. R90C showed significantly increased signalling activity compared to WT, and shifted to an alternative mechanism of activation that was independent of metalloprotease activity and dependent on TRPML, a calcium channel protein involved in endosomal/lysosomal fusion and fission and known to be involved in endosomal Notch activation pathways in *Drosophila*^15^. Signalling by WT, C455R and C212Y were less affected by TRPML inhibition than R90C, again pointing to distinct activation mechanisms for basal signalling. This shift in mechanistic requirements was correlated with an altered endocytic localisation with ECD removal likely enhanced in the EEA1 early endosome or occurring before reaching this location. The signalling activity of the C455R mutation is informative as it strongly reduces ligand-dependent induction of NOTCH3 signalling^23,30^. It is likely therefore that the basal signalling we observe is in large part due to a similar trans ligand-independent activation mechanism as has been observed for fly Notch^15^, which may arise from either ADAM10-dependent or independent mechanisms. This is consistent with previous work indicating that full-length wild type NOTCH3 has a basal level of constitutive activation^17,18^. We do not rule the involvement of cis-expressed ligands in the regulation of NOTCH3 endocytic trafficking and activation^44^, but it is noteworthy that the cis-interaction binding site of canonical Notch ligands is thought to occur through the same binding region as the trans-ligand interaction and is therefore also likely to be disrupted by the C455R mutation^45^.

The increased activity of CADASIL mutants, particularly R90C, is intriguing and is consistent with a number of recent reports which have suggested that CADASIL reflects a gain of function NOTCH3 activity. Increased NOTCH3 signalling has previously been identified in VSMC primary cultures derived from CADASIL patients^46^. CADASIL VSMCs were found to have increased cell proliferation, increased apoptosis and altered cytoskeletal morphology. NOTCH3 activation in the transgenic Notch3R169C mouse CADASIL model was also increased^25^. Increased activation of NOTCH3 in CADASIL was additionally observed in studies of VSMCs differentiated from induced pluripotent stem cells (iPSCs) generated from the fibroblasts of a CADASIL patient^47^. In another study, CADASIL mutant VSMCs, also differentiated from patient-derived iPSCs, were found to be more prone to apoptosis and failed to stabilise endothelial networks in a co-culture model, unlike wild-type VSMCs^42^. These defects were reversed by siRNA knockdown of NOTCH3, suggestive of a gain-of-function effect. Interestingly, in *Drosophila,* a number of Cysteine substitution mutants in EGF domains have been identified that display dominant, gain of function phenotypes^48^. Our work now offers a mechanistic explanation for increased Notch activity derived from EGF-module mutants that result in free cysteine mutants in Notch, through a ligand-independent activation pathway, involving shedding of the ECD during endocytic trafficking. We further suggest that subtle increases in this process over the long term may contribute to observed accumulation of ECD in CADASIL patients. The finding that R90C shifts its activation mechanism to a metalloprotease-independent mode indicates there is underlying heterogeneity in the ways the ECD can be detached from the full-length protein. This insight may offer routes to develop targeted therapies linked to specific mechanisms associated with particular mutants. Since R90C is amongst the most frequently occurring CADASIL mutants^33^ then identifying a targeted treatment for this mechanism would represent a significant step forward in the treatment of inherited vascular dementia. It remains to be seen whether other CADASIL mutants also switch to the alternative activation route. Even the mutants used in this study, which here activated like wild type, may switch to alternative mechanisms as the disease progresses or in response to additional environmental factors or other modifiers. For example, in *Drosophila*, switching between alternate metalloprotease-dependent and independent mechanisms can be brought about by changes in expression of ubiquitin ligase regulators that modify endosomal trafficking destinations of Notch^16^. Downregulating the function of ESCRTIII complex proteins, which regulate transfer of proteins between the endosomal membrane and the lumen, also activates both *Drosophila* Notch and WT human NOTCH3 in a metalloprotease-independent manner^16^.

There have been few studies that have addressed how heterogeneity of patient outcomes might be linked to differently located mutants. Recent data has suggested that CADASIL mutants in the N-terminal region have earlier onset of disease symptoms than more distally located mutants^22^. Other studies involving genome sequencing of large cohorts of individuals have revealed that CADASIL-like mutations in NOTCH3 are more common in the asymptomatic population than has been previously thought^12^. Such mutations have been shown to be predisposing to spontaneous stroke^12^. Altered NOTCH3 activity has also been linked to age-dependent small vessel disease^9^. Finding ways to tune NOTCH3 levels through modulation of its endocytic regulation may provide route to therapies for these conditions that are better tolerated in the long term than treatments through direct Notch pathway inhibition that are being considered as cancer therapeutics.

## Methods

### DNA Plasmid Constructs

NOTCH3 expression vectors were pcDNA3.1-Notch (WT, R90C, C455R, and C212Y) (Wang et al., 2007), a kind gift from Tao Wang (University of Manchester, UK). For empty vector control pcDNA3.1 was used (Thermo Fisher Scientific, Waltham, MA, USA). FP1 and FP2-NOTCH3 expression vectors were constructed in RP-HygroCMV (VectorBuilder Inc, IL, USA). FP1-NOTCH3 has in frame inserts of mTagBFP2 and mEGFP after amino acid 39 and 2129, respectively. FP2-NOTCH3 has in frame insertions of mScarlet-I and mNeonGreen at the same locations. For luciferase assays CBF-luciferase Notch reporter plasmid, was a kind gift from Keith Brennan (University of Manchester, UK)^49^, for constitutive Renilla expression we used pRL-CMV (Promega, Wi, USA).

### Cell Culture

hTERT Human Retinal Pigmented Epithelium-1 (hTERT-RPE-1, Clontech, CA, USA) cells were maintained in Dulbecco’s Modified Eagles Medium combined with high concentration of glucose, amino acids and vitamins with F-12 (DMEM, Sigma-Aldrich, MO, USA) supplemented with 1% penicillin-streptomycin (Sigma-Aldrich, MO, USA), 1% L-glutamine (Gibco, MA, USA) and 5% felt bovine serum (Sigma-Aldrich, MO, USA), Cells were fed every 2/3 days with fresh medium and incubated at 37°C in a humidified atmosphere containing 5% CO2. Experiments were conducted using cells at passage 6-18. hTERT-RPE-1 cells were obtained from Prof. Woodman (University of Manchester).

Michigan Cancer Foundation-7 (MCF-7) cells, (HTB-22, ATCC, VA, USA) were maintained in DMEM with low glucose and L-Glutamine supplemented with 1% penicillin-streptomycin (Sigma-Aldrich, MO, USA) and 5% felt bovine serum (Sigma-Aldrich, MO, USA) in T75 culture flasks (Corning, NY, USA). Cells were fed every 2/3 days with fresh medium and incubated at 37°C in a humidified atmosphere containing 5% CO2. Experiments were conducted using passage 16-20 MCF-7 cells.

Human embryonic stem cell line used was (Man-13)^50^. hESCs maintained in mTeSRTM1 Basal Medium (#85851 STEMCELL Technologies, BC, Canada) supplemented with mTeSRTM1, 5X Supplement #85852 (STEMCELL Technologies, BC, Canada). Cells cultured on six-well plate (Corning, NY, USA) coated with 2ml supplemented mTeSRTM 1and 10ng Vitronectin (rhVTN-N, A14700, Life Technologies, Paisley, UK) and 10uM Rock Inhibitor (Y-27632, STEMCELL Technologies, BC, Canada) per well of a six-well plate (Corning, NY, USA). Cells were fed using fresh media every day.

Man-13 cells were differentiated into VSMCs using an 18-day differentiation protocol^41^. 6-8 clusters/cm2 of Man-13 cells were seeded onto a VTN-N, 6-well plate in E8 (Gibco, MA, USA) medium supplemented with Y-27632 ROCK Inhibitor (10uM) for 24hr. Man-13 cells culture media was then swapped to E6 (Sigma-Aldrich, MO, USA) supplemented with 10uM SB-431542 (Tocris, Bioscience, Abingdon, UK) and 10ng/ml FGF2 (Peprotech, NJ, USA). The medium was replaced every 24hr with the same supplement for 5 days. On day 5 cells were washed with PBS without Ca2+/Mg2+ and dissociated by 1ml 0.5% EDTA/PBS. Cells were centrifuged at 500rpm for 5 minutes and then cell pellet was resuspended in E6 medium and seeded onto VTN-N coated six-well plates at 0.5×104 cells. E6 medium was supplemented with 2ng/ml TGF-ß1 (Peprotech, NJ, USA) and 5ng/ml PDGF-ß (100-14B, Peprotech, NJ, USA). The medium was replaced every 24hr with the same supplement until day 18 of differentiation.

### Cell Transfection

Prior to transfection, cells were cultured and seeded on 0.01% poly L-lysine-coated coverslips in a twelve-well or six-well plates (Corning, NY, USA) as required for 24hr Cells at 37°C to reach 50% confluency. Cells were transfected with desired plasmid and Genejuice® Transfection Reagent (Merck Millipore, MA, USA) at 1:3 ratio. First, 3ul of Genejuice® transfection reagents were diluted in 100ul Opti-MEM (Thermo Fisher Scientific, MA, USA) per sample and incubated at room temperature for 5 minutes. In the next step, desired amount of DNA was added to the dilution and incubated at room temperature for 15-20s to form a transfection complex. Then cells were washed with Opti-MEM media and exposed to transfection complex with an additional 900ul Opti-MEM media per well in six-well plates (Corning, NY, USA), for 2hrs, and then media was changed into fresh culture media.

### Antibodies

Primary antibodies used were polyclonal sheep anti-NOTCH3 ECD (SF1559, R&D systems, Minneapolis, MN, USA, used 1:500), monoclonal mouse anti-NOTCH3 ECD (1E4, Merck Life Science, Gillingham, UK, used 1:500), monoclonal Rabbit anti-ICD (D11B8, Cell Signaling technology Inc, MA, USA, used 1:500), polyclonal Rabbit anti-ICD (ab23456, Abcam, Cambridge, UK, used 1:500), monoclonal rat anti-ICD (8G5, Cell Signaling technology Inc, MA, USA, used 1:500), monoclonal mouse anti-EEA1 (E9Q6G, Merck-Millipore, MA, USA, used 1:200), monoclonal mouse anti-CD63 (RFAC4, Merck Life Science, Gillingham, UK, used 1:500), monoclonal rabbit anti-LAMP1 (D2D11, Cell Signaling technology Inc, MA, USA, used 1:500), monoclonal mouse anti-KDEL (10C3, ENZO Life Sciences, NY, USA, used 1:500), polyclonal anti-Rab11a (2413S, Cell Signaling Technology Inc, MA, USA, used 1:50), mouse monoclonal anti α-smooth muscle actin (1A4, Abcam, Cambridge, UK, used 1:500), polyclonal rabbit anti-Calponin, CNN1 (ab46794, Abcam, Cambridge, UK, used 1:500), monoclonal anti-Oct4 (9B7, R&D systems, MN, USA, used 1:500), mouse monoclonal anti-SOX2 (245610, R&D systems, MN, USA, used 1:500). Fluorescent-conjugated secondary antibodies were obtained from Invitrogen (CA, USA, used 1:500).

### Immunofluorescent (IF) Staining

Cells were washed with 2ml PBS prior to fixation with 1ml 4% paraformaldehyde (PFA) (PierceTM, Cat. no. 28906, Thermo Fisher Scientific,) in PBS for 20min at room temperature, and then incubated with 0.25% NH4Cl (Sigma Aldrich, MO, USA) for blocking PFA residues. Cells were washed again with PBS and then, to permeabilise, cells were incubated with 1ml 0.1% Triton X-100 (T9284, Sigma-Aldrich, MO, USA) in PBS for 10min following 2x wash with PBS. Next cells were incubated with 1ml 5% NDS (normal donkey serum, Jackson ImmunoResearch, PA, USA) in PBS for 30min to block any non-specific binding. Cells were incubated with 500ul of diluted desire primary antibody in blocking solution for 1hr at room temperature and then washed three times with 1ml solution and then incubated with 500ul of diluted desire secondary antibody in solution for 45 minutes at room temperature.

### Antibody Uptake Assay

To label surface NOTCH3, the transfected cells were washed with 1ml chilled fresh culture media on ice and then incubated with anti-ECD antibodies for 15 minutes on ice to slow down NOTCH3 endocytosis. Then cells were washed with chilled culture media and the with 1ml warm fresh culture medium. Wells were then incubated at 37°C/5%CO2 for various chase times. Coverslips were collected at each time point, washed with 1ml 1x PBS at room temperature and fixed with 4%PFA in PBS for 20min at room temperature. Cells were then blocked with 5%NDS/PBS for 30 minutes and subsequently incubated with desired secondary antibodies (1:500) for 45 minutes in dark at room temperature. Cells were then rinsed with 1ml of 5%NDS/PBS twice followed by one wash with 1ml PBS. Stained cells were then mounted in Vectorshield with DAPI (Vector Laboratories, CA, USA) and left at 4°C overnight. To label endocytosed NOTCH3, after the chase period, cells were fixed/permeabilised, as described above, and treated with anti-NOTCH3 ICD and endosomal pathway markers (anti-CD63 or anti-EEA1) followed by secondary antibodies.

### Immunofluorescence Imaging

Images were captured using Volocity (Perkin Elmer, Beaconsfield, UK) with an Orca-ER digital camera (Hamamatsu Photonics, Hammamatsu city, Japan) mounted on a M2 fluorescent microscope (Carl Zeiss, Oberkochen, Germany). Deconvolution was performed with three nearest neighbours using Openlab (Improvision/Perkin Elmer, Beaconsfield, UK) and processed in Photoshop (Adobe, CA, USA). For quantitation of colocalization in hTERT-RPE1 cells, Z-section images were recorded at x100 and deconvolved using Openlab with 3 nearest neighbours to remove out of focus signal. An image plane was selected midway through the cell body at the plane of focus for the nucleus. Images were uploaded to ImageJ using FIJI^51,52^ and colocalization quantified as Pearson’s coefficients using the JACOP (Just another colocalisation Protocol) plug-in^53^. For quantification of colocalisation in cytoplasmic organelles in MCF7 cells, a mask was created around the perimeter of the nucleus to exclude nuclear-localised anti-ICD staining from the quantification. For quantification of NOTCH3 accumulation in the ER, ImageJ was used to calculate the %area of KDEL staining occupied by NOTCH3, and the % of total intensity of NOTCH3 staining which was coincident with KDEL staining regions. For calculation of ECD and ICD separation in the early endosomes, EEA1-positive organelles were scored for number of puncta present as full-length, ECD-only and ICD-only localisations.

### Luciferase Notch Reporter Assay

The Dual-Luciferase® Reporter (DLRTM, E1910, Promega, WI, USA) Assay System was used to measure NOTCH3 signalling activity. The DLRTM assay is comprised of a Notch Response Element (NRE) Firefly luciferase reporter (p10xRbpj-luc)^49^ to measure NOTCH3 signalling activity and a constitutively expressed pRL-CMV (Promega, WI, USA) construct as an internal control for cell transfection efficiency and cell viability. pRL-CMV is a Renilla luciferase-expression (Promega, WI, USA) vector under CMV early promoter providing constitutive transcription. p10xRBPJ-luc is a Firefly luciferase reporter vector which has 10 copies of the canonical CBF/RBPJ transcription factor binding sites upstream of the firefly luciferase cDNA that act as an NRE. Both reporter vectors were co-transfected with the NOTCH3 plasmid in cell culture and to generate stabilised luminescence signals for both vectors in the same samples. The ratio of the two signals provides quantification of NOTCH3 activity.

### Use of Inhibitors

For treatment with inhibitors, MCF7 or hTERT-RPE1 cells were cultured for 48 hours with 10mM inhibitor in 2 ml culture medium, which was replaced after the first 24 hours. Inhibitors used were DAPT (Cell Guidance Systems, MO, USA), ML-SI1 (Scientific Laboratory Supplies, Nottingham, UK), BB94 (Stratech Scientific, Ely, UK), RO4929097 (Selleckeham, TX, USA).

## Acknowledgments

We thank Sue Kimber for the Man-13 hESC cell line, valuable discussion, and use of facilities, and Nicola Bates for advice with hESC cell culture and differentiation protocols. We further thank Tao Wang, Keith Brennan and Pat Caswell for vectors and reagents. We thank Antony Adamson, Wei-Hsiang and the FBMH Gene editing core facility for assistance with generating the MCF7 N3-knock out line. We acknowledge funding from BBSRC grant no. BB/V014218/1.

