## Supplemental Figures S1-4 for "Heterogeneity of NOTCH3 activation mechanisms uncovers therapeutic potential for targeted therapies for CADASIL"

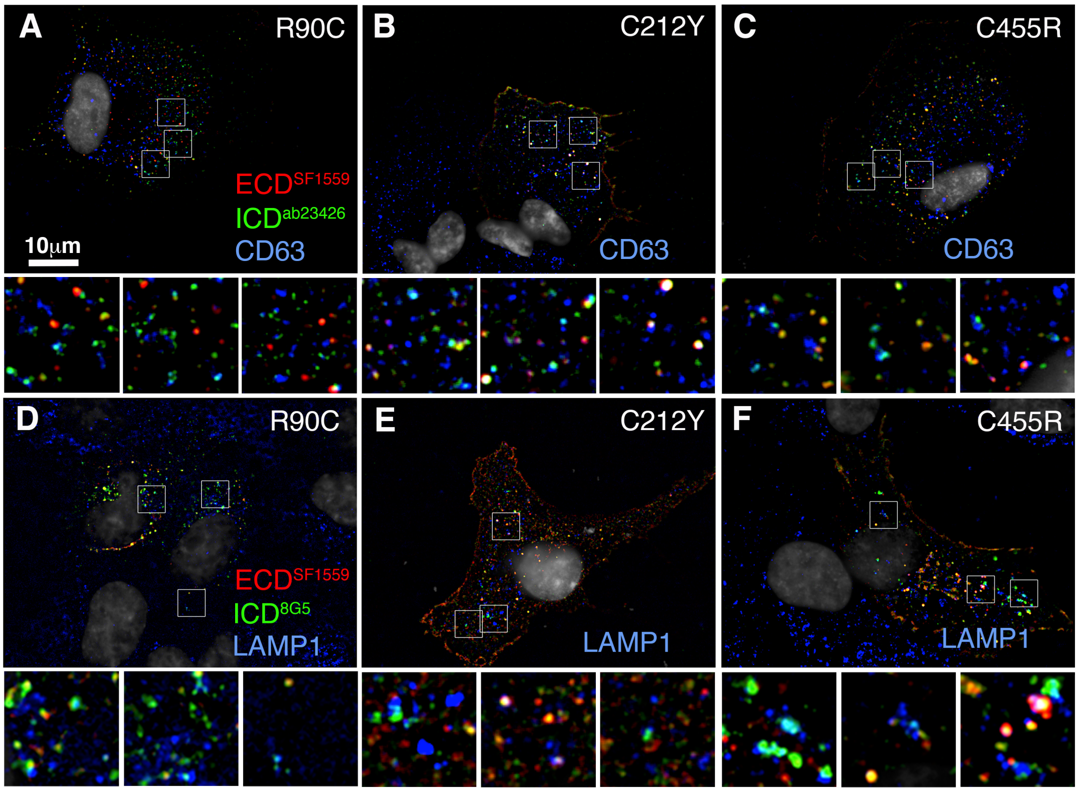


**Supplemental Figure S1. CADASIL mutant localisation in late endosomes and lysosomes.** **(A-C)** Localisation of ECD and ICD compared to CD63 late endosome, for R90C **(A)**, C212Y **(B)** and C455R **(C)**. Separated ICD puncta can be observed in late endosomes for all three mutants. **(D-F)** Similarly ICD-only puncta can be observed in LAMP1 positive lysosomes for R90C **(D)**, C212Y **(E)** and C455R **(F)**.

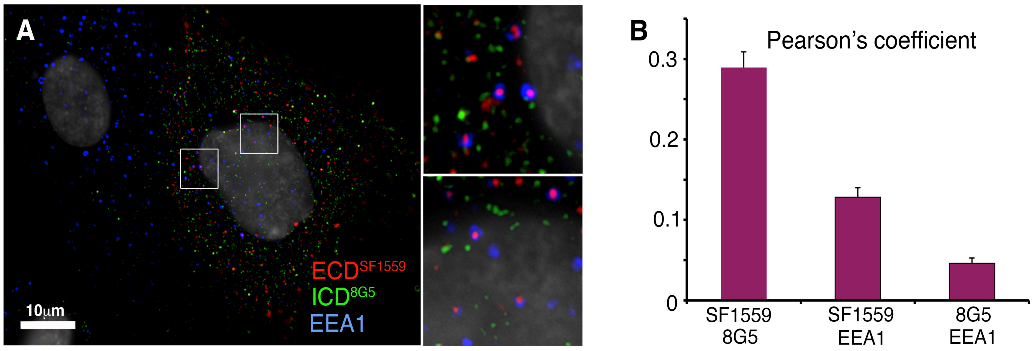


**Supplemental figure S2. Anti-ICD 8G5 epitope staining is excluded from early endosome location.**

**(A)** NOTCH3 expressing cell immunostained to compare the localisation of ECD and ICD-8G5 in EEA1-marked endosomes. ECD is clearly present in EEA1-endosomes, but there is little ICD-8G5 staining present in the endosome. Boxes show the regions enlarged in insets. **(B)** Quantification of the epitope colocalisations shown in A. Error bars are SEM, n=16.

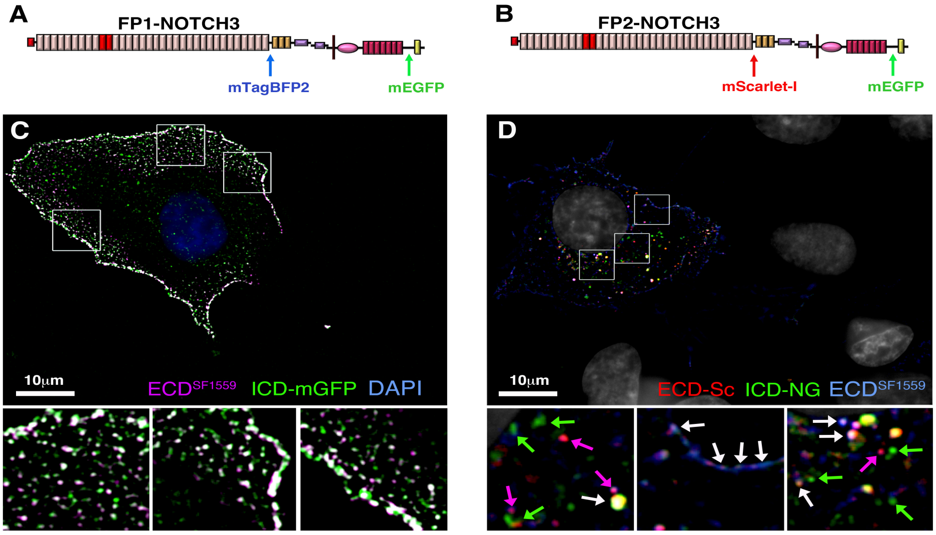


**Supplemental Figure S3. Fluorescent-tagged NOTCH3 shows ECD/ICD separation in intracellular compartments.**

**(A, B)** Schematic figure showing insertion sites of fluorescent-tagged proteins in WT NOTCH3 expression constructs FP1 and FP2. **(C)** Immunostaining of non-permeablised cells of surface-localised FP1-NOTCH3 with anti-ECD, which shows colocalisation in surface puncta with ICD-GFP, showing that full-length NOTCH3 is localised to the cell membrane. **(D)** Immunostaining of permeablised cells of FP2-NOTCH3. As well as full-length FP2-NOTCH3 (white arrows), separated mScarlet-I puncta colocalises with anti-ECD staining (magenta arrows) and ICD-GFP puncta can be seen which lack either mScarlet-I or anti-ECD staining (green arrows).

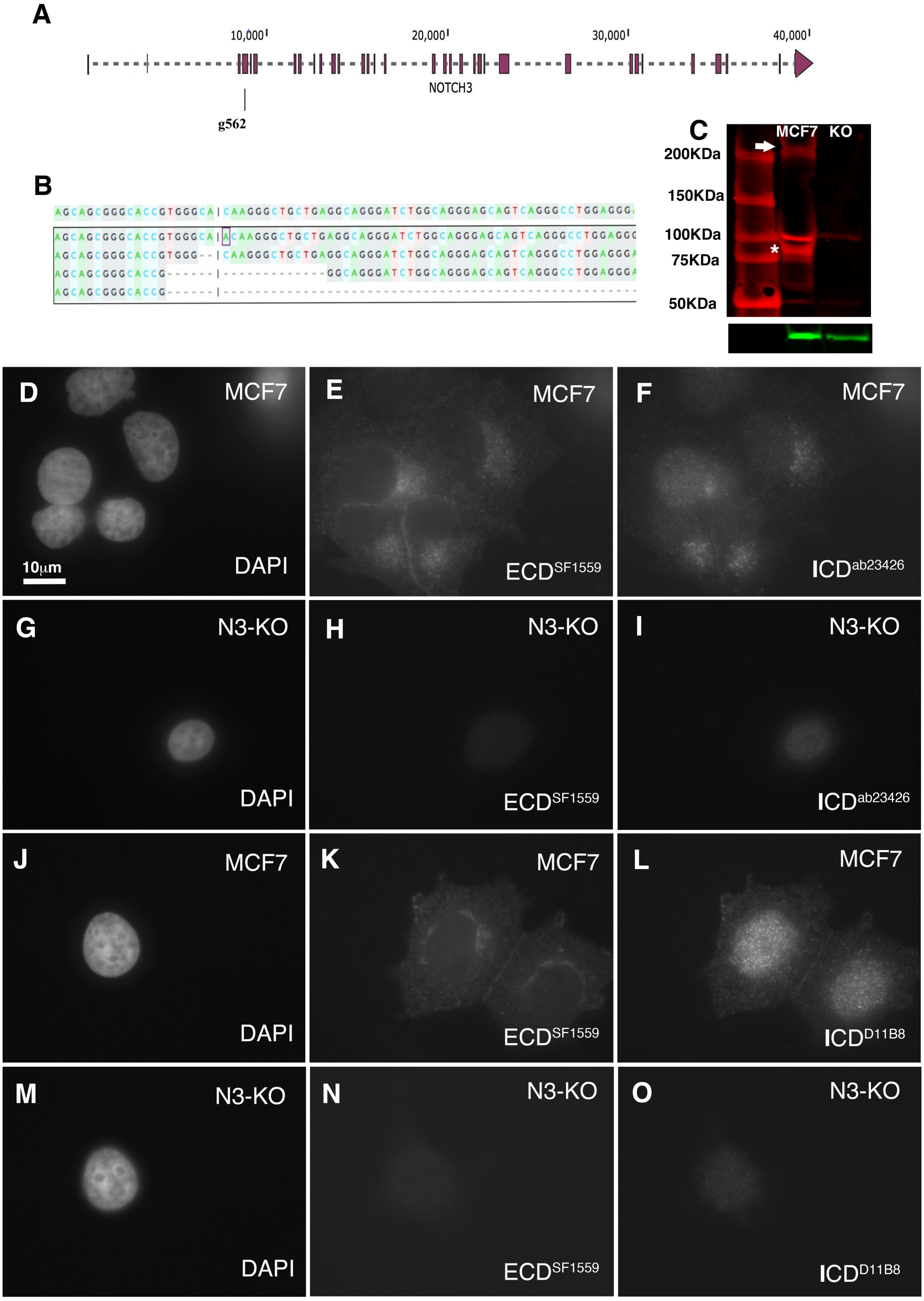


**Supplemental Figure S4. Validation of antibodies for immunolocalisation of endogenous NOTCH3.**

**(A)** NOTCH3 gene structure. Exons depicted by pink bars. The location of the guide RNA (g562), used for generating the CRISPR/Cas9 NOTCH3 knock out (N3-KO) is indicated. **(B)** After CRISPR, Sequencing of NOTCH3 genomic DNA, from line 2C7, in vicinity of guide site, revealed four sequences (boxed) containing deletions with frame shifts and identified no WT sequence. **(C)** Western blot stained with anti-ICD$^{D11B8}$ (red) showing loss of NOTCH3 expression in N3-KO cell line. Arrow indicates full-length NOTCH3 and asterisk, a membrane-tethered ICD. A band around 100kDa is likely a non-specific binding. **(D-O)** Immunofluresence staining of fixed and permeabilised parental MCF7 cells **(D-F, J-L)** and the 2C7 NOTCH3 knock-out line **(G-I, M-O)** for DAPI **(D, G, J, M)**, anti-ECD$^{SF1559}$ **(E, H, K, N)**, anti-ICD$^{ab23426}$ **(F, I)** and anti-ICD$^{D11B8}$ **(L, O)**. Antibody staining of parental MCF7 cells is distinctly above minimal background staining of ECD$^{SF1559}$ in NOTCH3-KO. While both ICD antibodies show some immunostaining of the nucleus in the NOTCH3-KO line, there is minimal staining in the cytoplasm compared to stronger staining in parental cells.
